# Spo0A suppresses *sin* locus expression in *Clostridioides difficile*

**DOI:** 10.1101/2020.02.28.968834

**Authors:** Babita Adhikari Dhungel, Revathi Govind

## Abstract

*Clostridioides difficile* is the leading cause of nosocomial infection and is the causative agent of antibiotic-associated diarrhea. The severity of the disease is directly associated with the production of toxins, and spores are responsible for the transmission and persistence of the organism. Previously we characterized *sin* locus regulators SinR and SinR’, where SinR is the regulator of toxin production and sporulation, while the SinR’ acting as its antagonist. In *Bacillus subtilis*, Spo0A, the master regulator of sporulation, regulates SinR, by regulating the expression of its antagonist *sinI*. However, the role of Spo0A in the expression of *sinR* and *sinR’* in *C. difficile* is not yet reported. In this study, we tested *spo0A* mutants in three different *C. difficile* strains R20291, UK1, and JIR8094, to understand the role of Spo0A in *sin* locus expression. Western blot analysis revealed that *spo0A* mutants had increased SinR levels. The qRT-PCR analysis for its expression further supported this data. By carrying out genetic and biochemical assays, we have shown that Spo0A can bind to the upstream region of this locus to regulates its expression. This study provides vital information that Spo0A regulates *sin* locus, which controls critical pathogenic traits such as sporulation, toxin production, and motility in *C. difficile*.

**IMPORTANCE:** *Clostridioides difficile* is the leading cause of antibiotic-associated diarrheal disease in the United States. During infection, *C. difficile spores* germinate, and the vegetative bacterial cells produce toxins that damage host tissue. In *C. difficile*, *sin* locus is known to regulate both sporulation and toxin production. In this study, we have shown that Spo0A, the master regulator of sporulation to control the *sin* locus expression. We performed various genetic and biochemical experiments to show that Spo0A directly regulates the expression of this locus by binding to its upstream DNA region. This observation adds new detail to the gene regulatory network that connects sporulation and toxin production in this pathogen.

## INTRODUCTION

*Clostridioides difficile* is a Gram-positive, anaerobic bacillus and is the principal causative agent of antibiotic-associated diarrhea and pseudomembranous colitis (1–3). Antibiotic use is the primary risk factor for the development of *C. difficile* associated disease because it disrupts normal protective gut flora and provides a favorable environment for *C. difficile* to colonize the colon. Two major pathogenic traits of *C. difficile* are toxin (toxin A and B) and spores (3–5). Deaths related to *C. difficile* increased by 400% between 2000 and 2007, in part because of the emergence of more aggressive *C. difficile* strains (6, 7). Robust sporulation and toxin production were suspected of contributing to the widespread of the *C. difficile* infections associated with these highly virulent strains (8–14). How *C. difficile* triggers toxin production and sporulation in the intestinal environment is only beginning to be understood.

Recently, we reported the identification and characterization of master regulator SinR in *C. difficile* that was found to regulate sporulation, toxin production, and motility (15). SinR in the Gram-positive model organism *B. subtilis* is well characterized and is known to regulate multiple pathways, including sporulation, competence, motility, and biofilm formation (16–18). In *B. subtilis*, SinR is encoded by the downstream gene of the two gene operon called *sin* (sporulation inhibition) locus and its transcription is driven from two promoters. The second gene in the operon, *sinR*, is transcribed by an internal promoter and is constitutively expressed. SinR represses the first committed (stage II) genes in the sporulation pathway (17).The promoter upstream of the operon is activated by phosphorylated Spo0A, leading to the expression of *sinI*, along with *sinR*. The SinI protein binds and inhibits the DNA binding activity of SinR (19–21). The combined effect of positive regulation by Spo0A∼P and the inactivation of the negative regulator SinR activates the sporulation pathway. In *C. difficile* also the *sin* locus is a two-gene operon and encodes for SinR and SinR’. In our initial characterization of the *C. difficile sin* locus, we have shown that disruption of *sin* locus (absence of both SinR and SinR’) resulted in asporogenic, less toxic, and less motile phenotype (15). Another study which reports that

*C. difficile sin* locus suppresses biofilm formation corroborates our finding (22). Further investigation showed that among the two regulators, SinR positively influences sporulation, toxin production, and motility, while SinR’ acts as an antagonist to SinR and control its activity. Since *sin* locus has a role in regulating various pathogenic traits in *C. difficile*, understanding the regulation of its expression is important. Earlier, we have shown that disrupting the first gene *sinR* in the operon to affects both the *sinR* and the downstream *sinR’* transcription. This led to the assumption that unlike *B. subtilis*, the *C. difficile sin* locus is transcribed from a single upstream promoter. Real-time RT PCR analysis of the cells grown *in vitro*, showed *sin* locus expression at 10-12 hour time point, indicating its tight regulation (15). From various gene expression data, we can observe that mutations in *sigH* and *spo0A* positively influence the expression of *sinRR’* (23–26) and mutations in *tcdR* to downregulate their expression (27). We have also demonstrated that CodY can directly bind to *sin* locus upstream DNA to transcriptionally repress its expression (15). In the same line of investigation, in this study, we discovered that Spo0A, the sporulation master regulator, represses *sinR* expression. The effect was directly caused by the specific binding of Spo0A to the promoter region upstream of the locus.

## MATERIAL AND METHODS

### Bacterial strains and growth conditions

*C. difficile* strains (Table S1) were grown in TY (Tryptose and Yeast extract) agar or broth culture in an anaerobic chamber which is maintained at 10% H_2_, 10% CO_2_ and 80% N_2_ as described previously (27–30). Lincomycin (Linc 20ug/ml) and thiamphenicol (Thio; 15 ug/ml) were added to the culture medium when required. S17-1, an *E.coli* strain used for conjugation (31), was cultured aerobically in LB (Luria-Bertani) broth or agar and was supplemented with ampicillin (100 µg/ml) or chloramphenicol (25 µg/ml) when necessary.

### General DNA techniques

Chromosomal DNA was extracted from *C. difficile* cultures using DNeasy Blood and Tissue Kit (Qiagen). PCR reactions were carried out using gene specific primers (Table S2). PCR products were extracted from the gel using Geneclean Kit (mpbio). Plasmid DNA was extracted using QIAprep Spin Miniprep Kit (Qiagen). Standard procedures were used to perform routine cloning.

### Construction and complementation of *C. difficile spo0A* mutant strains

The *spo0A* mutants in JIR8094 and UK1 strains were created using ClosTron gene knockout system as described previously (24, 25, 32, 33). Briefly, for *spo0A* disruption, the group II intron insertion site between nucleotides 178 and 179 in *spo0A* gene in the antisense orientation was selected using a web-based design tool called the Perutka algorithm. The designed retargeted intron was cloned into pMTL007-CE5 as described previously (34). The resulting plasmid pMTL007-CE5::*spo0A*-178-179a was transferred into *C*. *difficile* UK1 and JIR8094 cells by conjugation. The potential Ll.ltrB insertions within the target genes in the *C*. *difficile* chromosome was conferred by the selection of lincomycin resistant transconjugants in 20 μg/ ml lincomycin plates. PCR using gene-specific primers (Table S2) in combination with the EBS-U universal was performed to identify putative *C*. *difficile* mutants. *C. difficile spo0A* mutants were complemented by introducing pRG312 which contains the *spo0A* gene with the 300 bp upstream region, through conjugation. Complementation was confirmed by PCR and western blot analysis.

### Western blot analysis

*C. difficile* cultures for western blot analysis were harvested and washed in 1xPBS solution. The pellets were resuspended in sample buffer (Tris 80 mM; SDS 2%; Glycerol 10%) and lysed by sonication. The whole cell extracts were then centrifuged at 17,000 g at 4°C for 1 min. The lysate was heated at 100°C for 7 min and the proteins were separated by SDS-PAGE and electro-blotted onto PVDF membrane. The blots were then probed with specific primary and the secondary antibodies at a dilution of 1:10,000. Immuno-detection of proteins was done using ECL kit (Millipore) following the manufacturer’s recommendations and were developed using G-Box iChemi XR scanner. Blot images were overlapped with the original images of the membrane to visualize pre-stained marker.

### Construction of reporter plasmids and beta-glucuronidase assay

The *sin* locus upstream DNA regions of various lengths were amplified by PCR using specific primers with *KpnI* and *SacI* (Table S2) recognition sequences. R20291 strain chromosomal DNA was used as a template for this amplification. Plasmid pRPF185 carries a *gusA* gene for beta-glucuronidase under tetracycline-inducible (*tet*) promoter. The *tet* promoter was removed using *KpnI* and *SacI* digestion and was replaced with the *sin* locus upstream regions of various lengths to create plasmids pBA009, pBA029, pBA037, pBA038 and pBA039 (Table S1). The control plasmid pBA040 with promoter less *gusA* was created by digesting with *KpnI* and *SacI* to remove *tet* promoter and then self-ligated after creating blunt ends. Plasmids were introduced into R20291 and R20291::*spo0A* strains through conjugation as described previously (15, 27). The transconjugants were grown in TY medium in the presence of thiamphenicol (15 ug/ml) overnight. Overnight cultures were used as an inoculum at a 1:100 dilution to start a new culture. Bacterial cultures were harvested at 10 hr of growth and the amount of for beta-glucuronidase activity was assessed as described elsewhere (35, 36). Briefly, the cells were washed and resuspended in 1 ml of Z buffer (60 mM Na_2_HPO_4_.7H_2_O pH 7.0, 40 mM NaH_2_PO_4_.H_2_O, 10 mM KCl, 1mM MgSO_4_.7H_2_O and 50mM 2ME) and lysed by homogenization. The lysate was mixed with 160 ul of 6mM p-nitrophenyl β-D-glucuronide (Sigma) and incubated at 37°C. The reaction was stopped by the addition of 0.4 ml of 1.0 M NaCO_3_. β-Glucuronidase activity was calculated as descried earlier (35, 36).

### Mutagenesis of *sin* locus promoter region

Quick Change Lightning Site-Directed Mutagenesis Kit (Agilent Technologies) was used to carry out site directed mutagenesis whereby G and C residues of the potential Spo0A binding ‘0A’ boxes were substituted with A residues. The mutagenic oligonucleotide primers used are listed in Table S2.

### DNA binding

DNA binding was carried out as described elsewhere (37). Briefly, the promoter region of interest was biotin labelled and was coupled to immobilized Monomeric Avidin Resin (G Biosciences) in B/W Buffer (37). The DNA and the beads were incubated at room temperature for 30 min in a rotor. The bead-DNA complex was washed with TE Buffer to remove any unbound DNA. To prepare cell lysates, *C. difficile* R20291 strain was grown to late exponential phase (16 hour) in 500 ml TY medium, pH 7.4. After washing with 1XPBS, the cells were resuspended in BS/THES buffer (37) and lysed using French press. The whole lysate was centrifuged at 20,000 g for 30 min at 4°C and the supernatant was incubated with the bead-DNA complex and allowed to rotate at 4°C overnight. The bead-DNA-protein complex was washed with BS/THES Buffer (5 times). Elution was carried out with 50mM, 100 mM, and 200 mM NaCl in Tris-HCl pH 7.4. The eluates were analyzed by SDS-PAGE and Western Blotting using Spo0A specific antibody.

## RESULTS

### Elevated level of SinR is present in *spo0A* mutant

In *C. difficile*, we have previously shown that *sin locus* mutant is asporogenic and this phenotype is associated with downregulation of *spo0A* expression. Interestingly, disruption of *sinR’*, the second gene in the locus resulted in elevated levels of sporulation. This result suggested that SinR as a positive regulator of the sporulation. Gene expression data of *spo0A* mutants from different studies have shown an elevated levels of *sin locus* expression when compared to their respective parents (25, 32, 38). These observations taken together suggest that these two master regulators, Spo0A and SinR regulate each other’s transcription. To understand the possible regulatory relationship between SinR and Spo0A in *C. difficile*, we created *spo0A* mutant in two different *C. difficile* strains, JIR8094 and UK1 strain using the Clostron mutagenesis technique. Mutation in *spo0A* was confirmed by PCR (Fig. S1ABC) and western blot analysis using Spo0A specific antibodies (Fig. S1D). The *spo0A* mutant in R20291 obtained from Dena Lyras Lab (39) was also included in the analysis. As previously reported, mutation in *spo0A* resulted in the asporogenic phenotype (24, 25, 33, 39). For complementation, plasmid pRG312 carrying *spo0A* under its own promoter was into the mutants. Introduction of the *spo0A* expressing plasmid was successful in the JIR8094::*spo0A* and in R20291::*spo0A* mutants, but unsuccessful in the UK1::*spo0A* mutant. Heat resistant spores were observed in the complemented strains, however the levels were significantly lower than wild-type (Fig. S2). To test whether Spo0A influences expression of the *sin* locus genes, we performed quantitative reverse transcription-PCR (qRT-PCR) analysis of the *sinR* and *sinR’* transcripts in *spo0A* mutants and their respective parent strains. As previously reported, the level of *sinR* and *sinR’* transcripts were increased several folds, (Fig. 1A) in all three *spo0A* mutants compared to their parent strains. To further confirm this result, we performed western blot analysis using SinR specific antibodies. We grew the mutants and the respective parent strains in TY medium for 10 hours and observed the levels of SinR in their cytosol. We found that *spo0A* mutants in all three strains produced higher amounts of SinR compared to their respective parents (Fig. 1B, Fig.S1D). However, in our complementation of R20291::*spo0A* and JIR8094::*spo0A*, we did not see lower levels of SinR (Fig. 1B).

**Figure 1.**
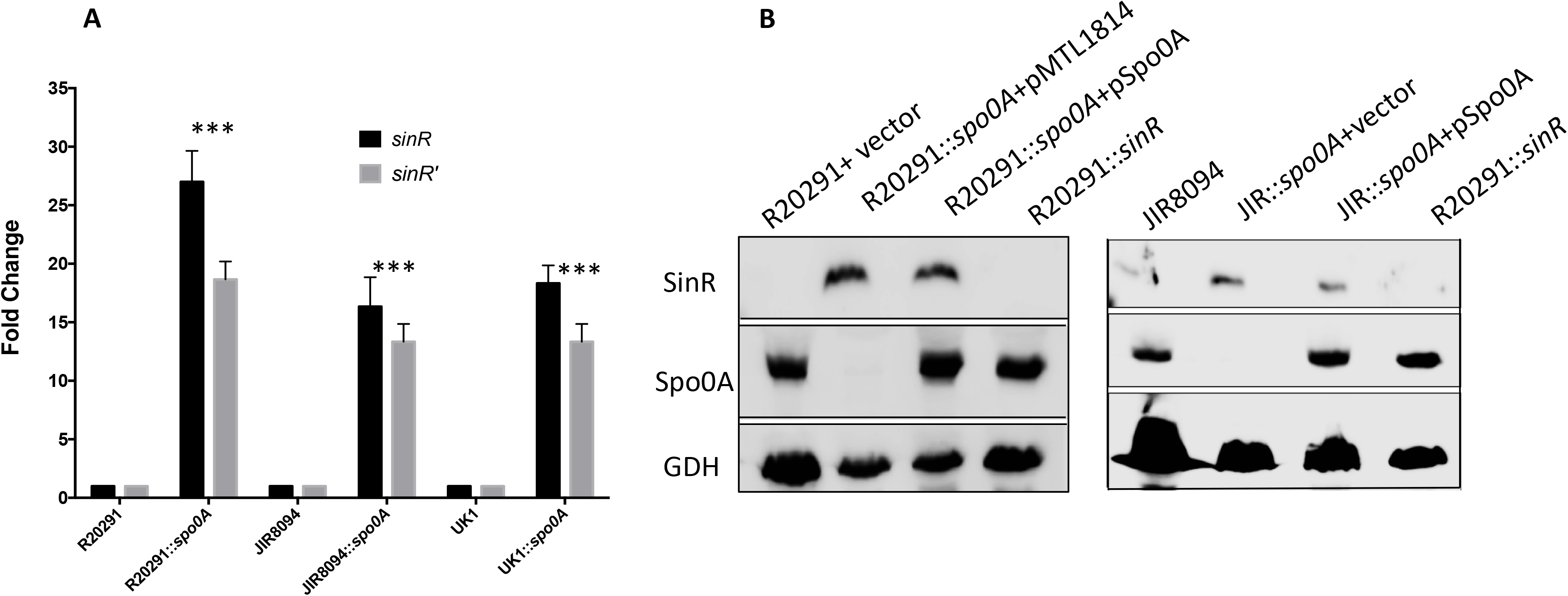
In the absence of Spo0A, *C. difficile* produces elevated levels of SinR. **A.** QRT-PCR results of *sin* locus transcripts in R20291; R20291::*spo0A*; JIR8094 and JIR8094::*spo0A* strains collected at 10 hr time point. The representative results from three independent experiments are shown. The asterisks (***) indicate statistical difference at a *P* value of <0.005. **B.** Western blot analysis parents (R20291 and JIR8094), and their respective *spo0A* mutants using SinR and Spo0A specific antibodies demonstrating upregulated SinR in the absence of Spo0A. The SinR negative R20291::*sinRR’* mutant served as a negative control. GDH detection using anti-GDH antibodies was used as loading control.

### Spo0A represses the expression of *sinR*

Spo0A is a transcriptional regulator and is a DNA binding protein. Spo0A binds to specific DNA sequences in the promoter region of its target gene to regulate their expression. To determine if the elevated levels of SinR observed in *spo0A* mutants is due to the repressor activity of Spo0A, we performed reporter fusion assays. We fused 600 bps of *sin* locus upstream DNA with the *gusA* reporter gene coding for beta-glucuronidase and the construct was introduced into the R2091::*spo0A* mutant and its parent strain. The plasmid carrying a promoter less *gusA* was used as a negative control. We also cloned the promoter region of *spoIIAB* known to be regulated by Spo0A, with the *gusA* and used this construct as a positive control. The *spoIIAB* promoter is positively regulated by Spo0A and was found to be active only in the parent strain and not in the *spo0A* mutant (Fig. 2A). We observed significantly higher beta glucuronidase activity when it was expressed from the *sin* locus promoter in the R20291::*spo0A* mutant strain compared to the parent strain, where very minimal reporter activity was recorded. This observation is consistent with our western blot results, where we detected elevated levels of SinR in *spo0A* mutant strains. Taken together these results suggest that Spo0A represses the transcription of *sinR* either directly or indirectly. To narrow down the Spo0A controlled region in the *sin* locus promoter, we cloned the 475 and 340 bps of the upstream DNA with the *gusA* gene and performed the reporter fusion assays. The reporter gene activity was similar in the cultures carrying the 600 bp as well as 340 bp upstream fusions (Fig. 2B). This indicated that both the *sin* locus promoter and the Spo0A regulated regions are present within this 340 bp region.

**Figure 2.**
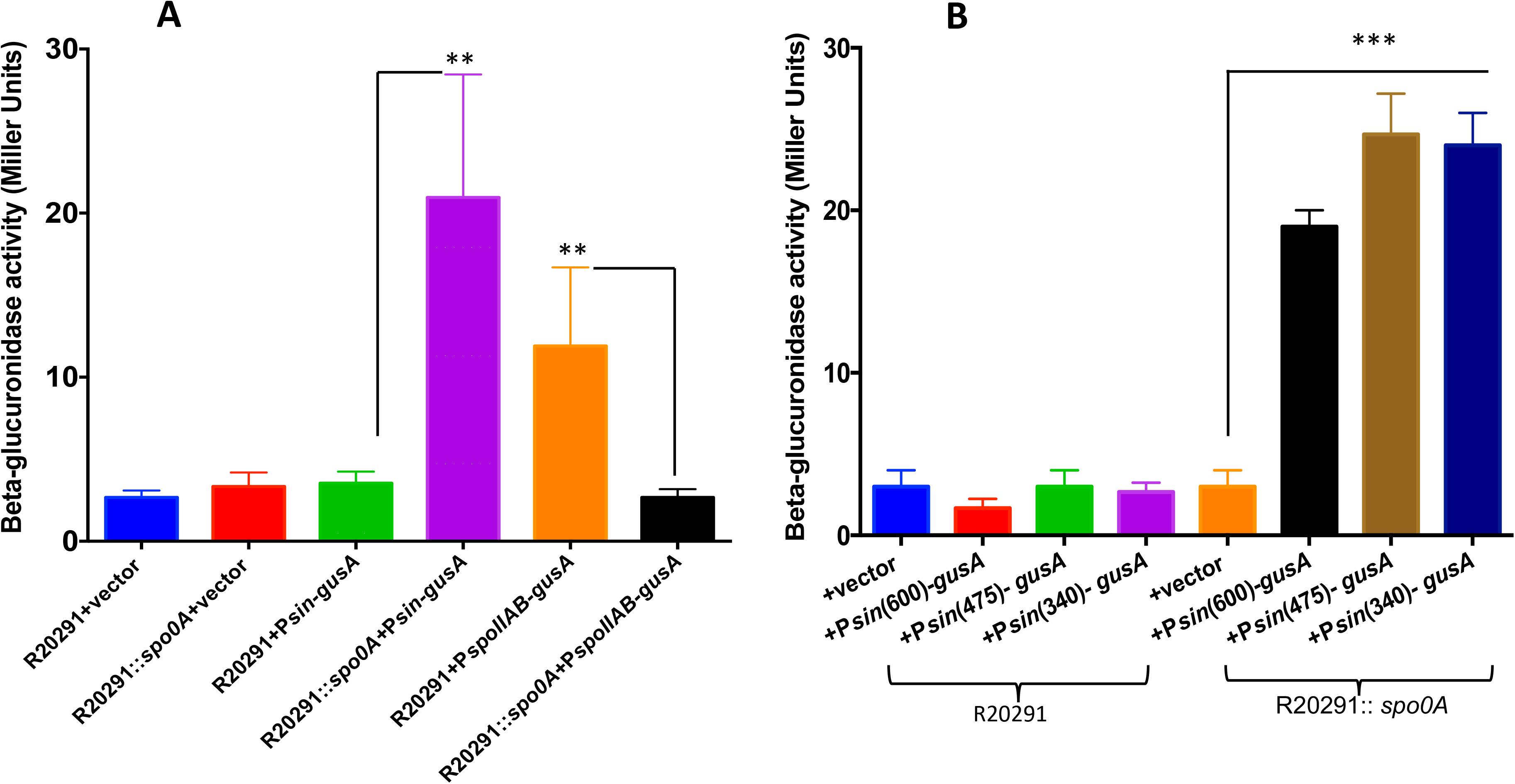
Spo0A represses *sin* locus expression. **A.** Beta-glucuronidase activity of the P*sin*-*gusA* fusions in the parent R20291 and R20291::*spo0A* mutant. Plasmid (pBA038) with *gusA* as the reporter gene fused to 600 bp of *sin* locus upstream. Plasmids carrying P*spoIIAB-gusA* (pBA029) and a promoter less *gusA* (pBA040) were used as positive and negative controls, respectively. **B.** Expression of beta-glucuronidase in parent R20291 and *spo0A* mutant carrying plasmids pBA037 (475 bps P*sin-gusA*), pBA038 (600 bp P*sin-gusA*) and pBA009 (340 bp P*sin-gusA*). The error bars in panels **A** and **B** correspond to standard errors of the means of results from 3 biological replicates, where ** and *** indicates P<0.05 and P<0.005, respectively (by two-tailed Student *t* test). At least three independent experiments were performed.

### Spo0A binds to the promoter region of *sinR*

The results in Figure 2AB show that expression of P*sin-gusA* was less in the R20291 background while the expression of the reporter gene was at higher levels in R20291::*spo0A* background. To determine whether the repression of *sinR* by Spo0A is due to Spo0A binding specifically to the promoter region of *sinR*, we carried out a DNA binding experiment. Considering that Spo0A needs to be phosphorylated to bind to the target DNA and inability to purify Spo0A-P, we did not attempt the *in vitro* electrophoretic gel shift assay. Instead, we used a biotin labelled DNA pulldown assay to determine the DNA binding ability of Spo0A under native conditions. The DNA segment representing the promoter region of *sinR* was biotinylated and was coupled to Immobilized Monomeric Avidin Resin. This bead-DNA complex was incubated with the cell lysate from the parent R20291 strain. The bound proteins were eluted and were run in SDS page and immunoblotted with Spo0A antibody. We first standardized the binding experiment by using *spo0IIAB* promoter region as a positive control. Spo0A protein could be detected in the elutes when the *spo0IIAB* upstream DNA was used the bait. The biotinylated *gluD* upstream DNA also processed similarly and served as a negative control. We applied the same protocol using the biotinylated 340 bp *sin* upstream DNA as bait. Results showed that it could pull down Spo0A, suggesting that Spo0A binds specifically to the promoter region of the *sin* locus (Fig. 3AB). Next, to narrow down the Spo0A binding site within that 340 bps we created three biotin-labelled fragments covering the first 118 bps (340-222 bps upstream), the last 140 bps and overlapping 135 bps mid-region (237 to 102 bps upstream) (Fig. 3A) and used them as bait in the pull-down experiment. More Spo0A was detected when the 140 bps mid-region was used as a bait compared to the first 118 bps (Fig. 3C). The Spo0A protein appears to bind with greater affinity to the middle fragment, when compared to the other two fragments. Since the biotin DNA pulldown assay is semi quantitative in nature, this assumption needs further validation. Spo0A could not be recovered from the elute from the binding of *gluD* upstream region, suggesting the specificity of the Spo0A binding with the *sin* locus promoter.

**Figure 3.**
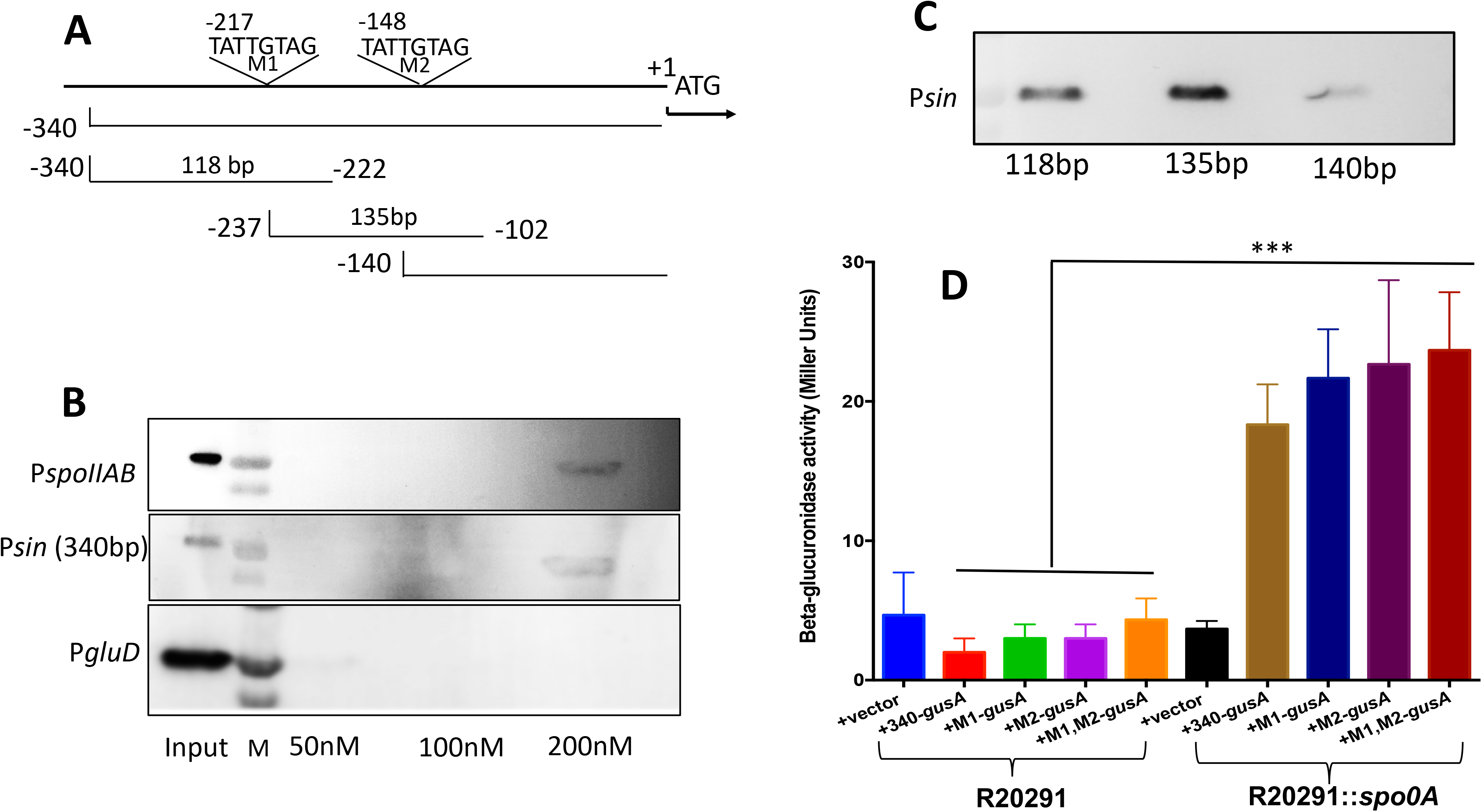
A. Spo0A binds to *sin* locus upstream DNA. Schematics of the 340 bp upstream *sin* locus denoting the general location of the TATTGTAG repeats (not to scale) with respect to the translation start (+1) of *sinR*. The lower lines indicate the location and the sizes of the DNA fragments used for the biotinylated DNA-pulldown assay. **B.** Western blot analysis using Spo0A specific antibody to detect endogenous Spo0A in input and elute fractions. For the biotin-DNA pulldown assay, the promoter regions of *spoIIAB* and *gluD* were used as positive and negative controls, respectively. **C.** Three DNA fragments (118 bp; 135 bp; 140 bp) spanning different regions of the 340 bp *sin* locus upstream were used independently to carry out the binding and Spo0A was detected as in panel B. **D.** Expression of beta glucuronidase in parent R20291 and R20291::*spo0A* mutant strains carrying plasmids with *gusA* as the reporter gene fused to the promoter of *sinR*. The predicted Spo0A box was mutagenized either on M1 or M2 site or both M1, M2. Strain carrying a promoter less *gusA* plasmid (pBA040) was used as control. Data represent the means ± standard errors of the means (SEM) (*n* = 3). The asterisks (***) in panel A indicate statistical difference at a *P* value of <0.005.

### Mutational analysis of the *sinR* upstream region

In *B. subtilis* Spo0A-P is known to bind to 7 bp DNA element 5’-TGNCGAA-3’, commonly known as Spo0A box (40). However, there are certain exceptions where Spo0A binds to degenerated Spo0A boxes with mismatches in the upstream of some targets (41, 42). The DNA binding domain of *C. difficile* Spo0A is highly homologous to *B. subtilis* Spo0A and the key residues of Spo0A known to mediate the interaction with the bases of the 0A box are highly conserved in *Bacillus* and *Clostridium* species (24, 43). In *C. difficile*, Spo0A is known to bind to *spo0A* upstream and *sigH* upstream. Both of these genes have the TGTCGAA consensus Spo0A box sequence (23, 44). *C. difficile* Spo0A also bind to the upstream of *spoIIAA*-*spoIIE*-*spoIIGA* operon with low affinity, where the binding sequence is a degenerated Spo0A box with TACGACA sequence (23). We scanned the upstream region of *sinR* for potential Spo0A binding consensus sequence. From the DNA binding pull-down experiment we could predict that Spo0A binds to sequences within the 237-102 bps upstream of *sin* locus. Classical Spo0A binding boxes couldn’t be identified in this region. However, two repeats with TATTGTAG sequences could be seen in this region. We mutated the TATTGTAG into TATTATAA, created reporter fusions with mutation in these sequences and analyzed their effect on the reported expression. None of the mutations affected the expression of the reporter gene. Mutations were introduced in the repeat sequence TAGTCTAT that occurred within the first 88 bps of *sin* locus upstream (data not shown). These changes also didn’t affect the *sin* locus expression. Even though, we could not identify the specific Spo0A binding boxes in the *sin* locus upstream, we have mapped the region to which it binds.

## DISCUSSION

Sporulation in a cell is an intense response to stress and is particularly expensive, in terms of both time and materials (45). The exact conditions and timing for sporulation are likely to be under strong selective pressure as both premature and belated spore production can have disastrous effects on cell growth and survival. In *B. subtilis*, the *sin* (*s*porulation *in*hibition) operon is central to the timing and early dynamics of this network (46–48), and its regulation is controlled by the sporulation master regulator Spo0A itself. In this study, we have demonstrated that similar to *B. subtilis*, the *sin* locus in *C. difficile* is also regulated by Spo0A. We created *spo0A* mutant in two different *C. difficile* strains JIR8094 and UK1, and included the previously created R20291::*spo0A* mutant in the analysis. As previously noted, we found both the JIR8094::*spo0A* and the UK1::*spo0A* strain to be asporogenic in nature. Complementation of sporulation phenotype was not complete in JIR8094::*spo0A* and we couldn’t complement the UK1::*spo0A* mutant strain even after multiple attempts for reasons unknown. Western blot analysis detected higher amount of SinR in all three different *spo0A* mutant strains. However, similar to the sporulation phenotype complementation of R20291::*spo0A* and JIR8094::*spo0A* did not lower the level of SinR. Unlike *B. subtilis* SinR, *C. difficile* SinR acts as a positive regulator of sporulation (15). How SinR controls sporulation is yet to be characterized. As we noted previously, *sin* locus is expressed at a low level only at a short window of time at 10-12 h of growth (15), suggesting that the presence of these regulators in the right amount at a specific time point could be important to regulate sporulation initiation and progression. In *spo0A* mutant however, *sinR* is expressed at a very high level which can affect the temporal nature of the sporulation pathway. It’s worth noting that asporogenic phenotype of *sinRR’* mutant could not be complemented either (15). Failures to complement a *spo0A* mutation have been previously observed in *C. difficile*. Two independent studies showed incomplete restoration of sporulation phenotype in R20291::*spo0A* (32, 39). However, when Deakin *et al.* (2012) tested the *spo0A* mutants in 630Δ*erm* and R20291 strains and they found *in vitro* levels of sporulation to be restored to wild type levels in their complemented derivative (33). When they tested the R20291 strains for toxin production, however, the complemented strain still produced increased levels of toxin compared to wild type (33). These observations suggest that this method of introducing *spo0A* using a multicopy plasmid may not be the suitable method for complementation considering the complex nature of Spo0A regulatory networks.

Like in many Gram-positive bacteria, Spo0A is the master regulator of sporulation in *C. difficile* (24, 33, 39). When the post-exponential phase begins, Spo0A activates the expression of the genes involved in the sporulation initiation process and positively regulates the sigma factor cascade required for sporulation (38). In many other pathogenic spore-forming bacteria the gene regulatory networks that influence sporulation and virulence are closely linked with each other (49–53). In *C. difficile*, the mutation in *spo0A* affected many pathogenic traits, including toxin production, flagella expression and biofilm formation (5, 32, 33, 39). Mackin *et. al* observed a clear increase in the production of toxin A and B upon disruption of *spo0A* in the ribotype 027 isolates R20291 and M7404 (39). In a similar study, Deakin *et al.* found that a R20291 *spo0A* mutant caused more severe disease in a murine model than the wild type strain, and associated this increase in severity with an increase in the amount of toxin A and toxin B produced by the mutant *in vitro* (33). Dawson et. al. showed that Spo0A in 630Δerm strain promotes sporulation cascade and biofilm formation in addition to negatively regulating virulence factor expression (toxins and flagella) (32). We found UK1::*spo0A* strain to produce higher toxins compared to its parent strain, while no significant difference was observed between the JIR8094 parent and JIR8094::*spo0A* mutant (Fig S3A). This observation was consistent with the previous report, where mutation in *spo0A* influenced the toxin production only in the 027 ribotype, which includes UK1 and R20291 strains, but not in the 630Δerm strain that belongs to 012 ribotype as the JIR8094. We measured the cytosolic toxins in all the three *spo0A* mutants and observed increased toxin production only in the R20291 and UK1 background, but not in the JIR8094 strain (Fig. S3A). Reduced biofilm formation was also found only in R20291::*spo0A*, UK1::*spo0A*, but not in JIR8094::*spo0A* strains (Fig. S3BC). The mechanism of Spo0A regulation over these pathways remains to be answered. In the *C. difficile* genome >100 open reading frames have potential 0A boxes within 500 bp of their start codons, indicating direct regulation by Spo0A (24). However, *tcdA* and *tcdB*, encoding toxin A and toxin B, respectively, are not among them, indicating the indirect influence of Spo0A on toxin production (24). Motility and biofilm formation could also be indirectly controlled by Spo0A, since many candidate regulators are encoded by the genes putatively under the direct control of Spo0A in *C. difficile* (24, 33). Our current finding of Spo0A mediated *sin* locus regulation can partly explain many of the phenotypes displayed by *spo0A* mutants, especially in the ribotype 027 strains (32, 39). In our initial characterization of the *sin* locus, we have shown decreased toxin production and motility in the absence of SinR and SinR’ (15). Expression of *sinR* alone was sufficient to complement these phenotypes and suggested SinR as a positive regulator of these pathways (15). We have further shown that SinR controls toxin production by regulating *sigD*, a sigma factor that positively regulates *tcdR*, which is needed for the transcription of toxin genes (15, 54, 55). SigD is also needed for the transcription of flagellar operon in *C. difficile* (54, 55). In this study, we have shown an increased SinR production in the absence of Spo0A (Fig. 2B). Quantitative RT-PCR results showed increased expression of *sigD*, *tcdR*, and *tcdB* in the R20291::*spo0A* and in UK1::*spo0A*, compared to their respective parent strains (Fig. S4). Increased *sigD* expression can lead to increased flagellar, toxin production and a reduced biofilm formation in the *spo0A* mutant (22, 32) (Fig. S3BC).

In this study, we have shown that Spo0A binds to the promoter of the *C. difficile sin* locus and suppresses the expression of both *sinR* and *sinR’*. Previously, we have shown that disrupting *sinR* by insertion mutagenesis affects both *sinR* and *sinR’* transcription (15), suggesting that *sinRR’* is transcribe as a bicistronic message. Our QRT-PCR analysis detected lower levels of *sinR’* transcripts than the *sinR* transcripts. Since the reduction is observed both in the parent and the *spo0A* mutants, we can conclude that this effect is independent of Spo0A. The reason for the reduced level of SinR’ over SinR is not clear. In *B*. *subtilis* the polycistronic *sinRI* transcripts are produced from two upstream promoters. The monocistronic *sinR* transcripts are driven from a promoter located within the coding region of *sinI*. In *B. subtilis*, Spo0A activates the expression of *sinI* by binding to the upstream promoter of the operon and indirectly regulating SinR activity (19, 20). Similar to *B. subtilis*, in *C. difficile* also there is a possibility that *sinR’* may have an independent promoter within the *sinR* coding sequence and could be controlled by an unknown regulator in *C. difficile*.

In summary, we have demonstrated that Spo0A, the master regulator of sporulation, regulates the expression of *sin* locus. We have further shown that Spo0A can bind to the upstream region of *sin* locus have successfully mapped the region to which it binds.

## Supporting Information

**S1 Fig. Construction and confirmation of the *spo0A* mutant in *C. diffcile* JIR8094 and UK1 strain. (A)** Schematic representation of ClostTron (group II intron) mediated insertional inactivation of *spo0A* gene in *C. difficile*. **(B)** PCR verification of the intron insertion and complementation of *spo0A* in JIR8094 with intron-specific primer EBS universal [EBS(U)] with *spo0A* specific primers ORG 551 and ORG 552. **(C)** PCR verification of the intron insertion in UK1 strain with EBS(U), ORG 551 and ORG 552. **(D)** Western blot analysis of Spo0A and SinR production in UK1 and UK1::*spo0A*. GDH was used as loading control.

**S2 Fig. Sporulation in *spo0A* mutants**. Percentage sporulation (the CFU/ml from ethanol resistant spores) of the parent and *spo0A* mutants. The representative results from three independent experiments are shown.

**S3 Fig**. **Toxin gene transcription and biofilm formation in R20291::*spo0A* mutant strain. (A)** Toxin production measured by ELISA. Statistical analysis was performed using student t-test. (**; <0.05 *P* value). **(B)** Crystal violet stained biofilm in the 12 wells tissue culture plate, showing poor biofilm formation in R20291::*spo0A* and in UK1::*spo0A* mutants.(**C**) Quantification of crystal violet dye attached to the cells forming biofilms.

**S4 Fig. qRT-PCR analysis.** Relative expression of the transcripts of *tcdR, sigD, and sinR* genes from *C*. *difficile* parent and *spo0A* mutant strains in JIR8094, R20291 *and* UK1 background. RNA was collected at 10 hr time point.

## REFERENCE

1. Guh AY, Kutty PK. 2018. Clostridioides difficile Infection. Ann Intern Med 169:ITC49–ITC64.

2. Prete RD, Ronga L, Addati G, Magrone R, Abbasciano A, Decimo M, Miragliotta G. 2019. Clostridium difficile. A review on an emerging infection. La Clinica Terapeutica 1:e41–e47.

3. Awad MM, Johanesen PA, Carter GP, Rose E, Lyras D. 2014. Clostridium difficile virulence factors: Insights into an anaerobic spore-forming pathogen. Gut Microbes 5:579–593.

4. Schäffler H, Breitrück A. 2018. Clostridium difficile – From Colonization to Infection. Front Microbiol 9.

5. Underwood S, Guan S, Vijayasubhash V, Baines SD, Graham L, Lewis RJ, Wilcox MH, Stephenson K. 2009. Characterization of the Sporulation Initiation Pathway of Clostridium difficile and Its Role in Toxin Production. Journal of Bacteriology 191:7296–7305.

6. 2016. CDC Press Releases. CDC. https://www.cdc.gov/media/releases/2015/p0225-clostridium-difficile.html

7. Lessa FC, Mu Y, Bamberg WM, Beldavs ZG, Dumyati GK, Dunn JR, Farley MM, Holzbauer SM, Meek JI, Phipps EC, Wilson LE, Winston LG, Cohen JA, Limbago BM, Fridkin SK, Gerding DN, McDonald LC. 2015. Burden of Clostridium difficile Infection in the United States. New England Journal of Medicine 372:825–834.

8. Depestel DD, Aronoff DM. 2013. Epidemiology of Clostridium difficile infection. J Pharm Pract 26:464–475.

9. Akerlund T, Persson I, Unemo M, Noren T, Svenungsson B, Wullt M, Burman LG. 2008. Increased Sporulation Rate of Epidemic Clostridium difficile Type 027/NAP1. Journal of Clinical Microbiology 46:1530–1533.

10. O’Connor JR, Johnson S, Gerding DN. 2009. Clostridium difficile Infection Caused by the Epidemic BI/NAP1/027 Strain. Gastroenterology 136:1913–1924.

11. Camacho-Ortiz A, López-Barrera D, Hernández-García R, Galván-De Los Santos AM, Flores-Treviño SM, Llaca-Díaz JM, Maldonado-Garza HJ, Garza HJM, Bosques-Padilla FJ, Garza-González E. 2015. First report of Clostridium difficile NAP1/027 in a Mexican hospital. PLoS ONE 10:e0122627.

12. Giancola SE, Williams RJ, Gentry CA. 2018. Prevalence of the Clostridium difficile BI/NAP1/027 strain across the United States Veterans Health Administration. Clin Microbiol Infect 24:877–881.

13. Fatima R, Aziz M. 2019. The Hypervirulent Strain of Clostridium Difficile: NAP1/B1/027 - A Brief Overview. Cureus 11.

14. Stabler RA, He M, Dawson L, Martin M, Valiente E, Corton C, Lawley TD, Sebaihia M, Quail MA, Rose G, Gerding DN, Gibert M, Popoff MR, Parkhill J, Dougan G, Wren BW. 2009. Comparative genome and phenotypic analysis of Clostridium difficile 027 strains provides insight into the evolution of a hypervirulent bacterium. Genome Biology 10:R102.

15. Girinathan BP, Ou J, Dupuy B, Govind R. 2018. Pleiotropic roles of Clostridium difficile sin locus. PLOS Pathogens 14:e1006940.

16. Kearns DB, Chu F, Branda SS, Kolter R, Losick R. 2005. A master regulator for biofilm formation by Bacillus subtilis. Molecular Microbiology 55:739–749.

17. Cervin MA, Lewis RJ, Brannigan JA, Spiegelman GB. 1998. The Bacillus subtilis regulator SinR inhibits spoIIG promoter transcription in vitro without displacing RNA polymerase. Nucleic Acids Res 26:3806–3812.

18. Mandic-Mulec I, Doukhan L, Smith I. 1995. The Bacillus subtilis SinR protein is a repressor of the key sporulation gene spo0A. J Bacteriol 177:4619–4627.

19. Bai U, Mandic-Mulec I, Smith I. 1993. SinI modulates the activity of SinR, a developmental switch protein of Bacillus subtilis, by protein-protein interaction. Genes Dev 7:139–148.

20. Milton ME, Draughn GL, Bobay BG, Stowe SD, Olson AL, Feldmann EA, Thompson RJ, Myers KH, Santoro MT, Kearns DB, Cavanagh J. 2019. The Solution Structures and Interaction of SinR and SinI: Elucidating the Mechanism of Action of the Master Regulator Switch for Biofilm Formation in Bacillus subtilis. Journal of Molecular Biology.

21. Shafikhani SH, Mandic-Mulec I, Strauch MA, Smith I, Leighton T. 2002. Postexponential Regulation of sin Operon Expression in Bacillus subtilis. Journal of Bacteriology 184:564–571.

22. Poquet I, Saujet L, Canette A, Monot M, Mihajlovic J, Ghigo J-M, Soutourina O, Briandet R, Martin-Verstraete I, Dupuy B. 2018. Clostridium difficile Biofilm: Remodeling Metabolism and Cell Surface to Build a Sparse and Heterogeneously Aggregated Architecture. Front Microbiol 9.

23. Saujet L, Monot M, Dupuy B, Soutourina O, Martin-Verstraete I. 2011. The Key Sigma Factor of Transition Phase, SigH, Controls Sporulation, Metabolism, and Virulence Factor Expression in Clostridium difficile. Journal of Bacteriology 193:3186–3196.

24. Rosenbusch KE, Bakker D, Kuijper EJ, Smits WK. 2012. C. difficile 630Derm Spo0A Regulates Sporulation, but Does Not Contribute to Toxin Production, by Direct High-Affinity Binding to Target DNA. PLOS ONE 7:12.

25. Pettit LJ, Browne HP, Yu L, Smits W, Fagan RP, Barquist L, Martin MJ, Goulding D, Duncan SH, Flint HJ, Dougan G, Choudhary JS, Lawley TD. 2014. Functional genomics reveals that Clostridium difficile Spo0A coordinates sporulation, virulence and metabolism. BMC Genomics 15:160.

26. Nawrocki KL, Edwards AN, Daou N, Bouillaut L, McBride SM. 2016. CodY-Dependent Regulation of Sporulation in Clostridium difficile. Journal of Bacteriology 198:2113–2130.

27. Girinathan BP, Monot M, Boyle D, McAllister KN, Sorg JA, Dupuy B, Govind R. 2017. Effect of tcdR Mutation on Sporulation in the Epidemic Clostridium difficile Strain R20291. mSphere 2.

28. Girinathan BP, Braun SE, Govind R. 2014. Clostridium difficile glutamate dehydrogenase is a secreted enzyme that confers resistance to H_2_O_2_. Microbiology 160:47–55.

29. Girinathan BP, Braun S, Sirigireddy AR, Lopez JE, Govind R. 2016. Importance of Glutamate Dehydrogenase (GDH) in Clostridium difficile Colonization In Vivo. PLOS ONE 11:e0160107.

30. Govind R, Dupuy B. 2012. Secretion of Clostridium difficile Toxins A and B Requires the Holin-like Protein TcdE. PLOS Pathogens 8:e1002727.

31. Teng F, Murray BE, Weinstock GM. 1998. Conjugal transfer of plasmid DNA from Escherichia coli to enterococci: a method to make insertion mutations. Plasmid 39:182–186.

32. Dawson LF, Valiente E, Faulds-Pain A, Donahue EH, Wren BW. 2012. Characterisation of Clostridium difficile biofilm formation, a role for Spo0A. PLoS ONE 7:e50527.

33. Deakin LJ, Clare S, Fagan RP, Dawson LF, Pickard DJ, West MR, Wren BW, Fairweather NF, Dougan G, Lawley TD. 2012. The Clostridium difficile spo0A gene is a persistence and transmission factor. Infect Immun 80:2704–2711.

34. Heap JT, Pennington OJ, Cartman ST, Carter GP, Minton NP. 2007. The ClosTron: A universal gene knock-out system for the genus Clostridium. Journal of Microbiological Methods 70:452–464.

35. Dupuy B, Sonenshein AL. 1998. Regulated transcription of Clostridium difficile toxin genes. Molecular Microbiology 27:107–120.

36. Mani N, Lyras D, Barroso L, Howarth P, Wilkins T, Rood JI, Sonenshein AL, Dupuy B. 2002. Environmental Response and Autoregulation of Clostridium difficile TxeR, a Sigma Factor for Toxin Gene Expression. Journal of Bacteriology 184:5971–5978.

37. Jutras BL, Verma A, Stevenson B. 2012. Identification of novel DNA-binding proteins using DNA-affinity chromatography/pull down. Curr Protoc Microbiol Chapter 1:Unit1F.1.

38. Fimlaid KA, Bond JP, Schutz KC, Putnam EE, Leung JM, Lawley TD, Shen A. 2013. Global analysis of the sporulation pathway of Clostridium difficile. PLoS Genet 9:e1003660.

39. Mackin KE, Carter GP, Howarth P, Rood JI, Lyras D. 2013. Spo0A Differentially Regulates Toxin Production in Evolutionarily Diverse Strains of Clostridium difficile. PLoS ONE 8:e79666.

40. Asayama M, Yamamoto A, Kobayashi Y. 1995. Dimer form of phosphorylated Spo0A, a transcriptional regulator, stimulates the spo0F transcription at the initiation of sporulation in Bacillus subtilis. J Mol Biol 250:11–23.

41. Baldus JM, Green BD, Youngman P, Moran CP. 1994. Phosphorylation of Bacillus subtilis transcription factor Spo0A stimulates transcription from the spoIIG promoter by enhancing binding to weak 0A boxes. J Bacteriol 176:296–306.

42. Chen G, Kumar A, Wyman TH, Moran CP. 2006. Spo0A-Dependent Activation of an Extended −10 Region Promoter in Bacillus subtilis. Journal of Bacteriology 188:1411–1418.

43. Molle V, Fujita M, Jensen ST, Eichenberger P, González-Pastor JE, Liu JS, Losick R. 2003. The Spo0A regulon of Bacillus subtilis. Mol Microbiol 50:1683–1701.

44. Saujet L, Pereira FC, Henriques AO, Martin-Verstraete I. 2014. The regulatory network controlling spore formation in Clostridium difficile. FEMS Microbiology Letters 358:1–10.

45. Hutchison EA, Miller DA, Angert ER. 2014. Sporulation in Bacteria: Beyond the Standard Model. Microbiol Spectr 2.

46. Grossman AD. 1995. Genetic Networks Controlling the Initiation of Sporulation and the Development of Genetic Competence in Bacillus Subtilis. Annual Review of Genetics 29:477–508.

47. Gaur NK, Dubnau E, Smith I. 1986. Characterization of a cloned Bacillus subtilis gene that inhibits sporulation in multiple copies. J Bacteriol 168:860–869.

48. Gaur NK, Cabane K, Smith I. 1988. Structure and expression of the Bacillus subtilis sin operon. J Bacteriol 170:1046–1053.

49. Dale JL, Raynor MJ, Ty MC, Hadjifrangiskou M, Koehler TM. 2018. A Dual Role for the Bacillus anthracis Master Virulence Regulator AtxA: Control of Sporulation and Anthrax Toxin Production. Front Microbiol 9:482.

50. Deng C, Peng Q, Song F, Lereclus D. 2014. Regulation of cry gene expression in Bacillus thuringiensis. Toxins (Basel) 6:2194–2209.

51. Dürre P. 2014. Physiology and Sporulation in Clostridium. Microbiology Spectrum 2.

52. Li J, Paredes-Sabja D, Sarker MR, McClane BA. 2016. Clostridium perfringens Sporulation and Sporulation-Associated Toxin Production. Microbiol Spectr 4.

53. Mi E, Li J, McClane BA. 2018. NanR Regulates Sporulation and Enterotoxin Production by Clostridium perfringens Type F Strain F4969. Infect Immun 86.

54. El Meouche I, Peltier J, Monot M, Soutourina O, Pestel-Caron M, Dupuy B, Pons J-L. 2013. Characterization of the SigD regulon of C. difficile and its positive control of toxin production through the regulation of tcdR. PLoS ONE 8:e83748.

55. McKee RW, Mangalea MR, Purcell EB, Borchardt EK, Tamayo R. 2013. The second messenger cyclic Di-GMP regulates Clostridium difficile toxin production by controlling expression of sigD. J Bacteriol 195:5174–5185.

